# The role of genomic vs. epigenomic variation in shaping patterns of convergent transcriptomic variation across continents in a young species complex

**DOI:** 10.1101/784231

**Authors:** Clément Rougeux, Martin Laporte, Pierre-Alexandre Gagnaire, Louis Bernatchez

## Abstract

Repeated adaptive divergence in replicates of phenotypic diversification offers a propitious context to identify the molecular bases associated to adaptive divergence. A currently hotly debated topic pertains to the relative role of genomic vs. epigenomic variation in shaping patterns of phenotypic variation at the gene expression level. Here, we combined genomic, epigenomic and transcriptomic information from 64 individuals in order to quantify the relative role of SNPs and DNA methylation variation in the repeated evolution of four limnetic-benthic whitefish species pairs from Europe and North America. We first found evidence for 149 convergent differentially methylated regions (DMRs) between species across continents, which significantly influenced levels of gene expression. Hyper-methylated DMRs in the limnetic species were globally associated to an expression repression relatively to benthic species, and inversely. Furthermore, we identified 108 convergent genetic variants (eQTLs) associated to gene expression differences between species. Gene expression differences were more pronounced in genes harbouring eQTL compared to those associated with DMRs, thus revealing a greater effect of eQTLs on gene expression. Multivariate analyses allowed partitioning the relative contribution of epi-/genomic changes and their association to gene expression variation. Most of the gene expression variation was significantly explained by genomic (4.1%) and putatively genomic-epigenomic interactive variation (46.7%), while “pure” epigenomic variation explained marginally 2.3% of the gene expression variation across continents. This study provides a rare qualitative and quantitative documentation of the relative role of genomic, DNA methylation and their interaction in shaping patterns of convergent gene expression during the process of ecological speciation.

## INTRODUCTION

Repeated adaptive divergence in replicates of phenotypic diversification offers a propitious context to identify the molecular bases and mechanisms associated to adaptive divergence. The expression of new phenotype may be influenced by the underlying genomic basis, the environmental conditions and their interaction on the considered traits. Genetic variation inducing repeated phenotypic diversification in independent diverging species pairs may i) originate from independent *de novo* mutation [1-3], ii) have a single origin and spread by gene flow [4-6], or iii) be independently recruited from standing genetic variation [7,8]. However, identical *de novo* mutations and gene flow are respectively unlikely to occur repeatedly among independent populations, and between geographically isolated populations. Moreover, maintaining standing genetic variation have been shown to increase the probability of genetic convergence in the evolution of complex traits (*i*.*e*., underlying a polygenic selection) [9,10], despite putative genetic redundancy.

The ability of organisms to rapidly diverge and adapt to new environmental conditions can be constrained by the genomic bases associated to phenotypic traits that will experience selection, as it might be eased by long term maintenance of genetic polymorphism from the ancestral genetic pool [8,11,12]. Such genetic polymorphism has been characterized to be heterogeneously distributed along the genome [13]. Indeed, most of the genetic variation occurs in inter-genic and regulatory regions, relatively to coding regions [14,15], although effect size of genetic variation in coding regions has been documented to contribute to gene expression variation similarly to non-coding genetic variants [16]. Such genetic polymorphism can act as *trans*- and/or *cis*-acting genetic variants [17] that might induce variation in gene expression regulation [18]. Moreover, *cis*-acting variants localized in coding sequences or into regulatory regions (non-coding gene sequence) might be inherited through generations [19], and have been described as a major source of phenotypic diversification via adaptive divergence in the early stages of ecological speciation [17,20-23]. Despite their direct effect on the gene expression regulation, other modifications and molecular mechanisms such as epigenetic variation can affect the transcriptional activity of related genes [24].

Epigenetics refers to changes in gene function without any alteration in gene sequence that are transmitted through mitosis as well as meiosis [25]. As such, epigenetics can modify the individual phenotype during development and/or in response to a changing environment without altering the DNA sequence [26]. Epigenetic variation has been categorized in three groups according to their level of independence with genetic variation, which are: i) ‘obligatory’ when relying completely on genetic variation, ii) ‘facilitated’ when indirectly potentiated by the genotype, and iii) ‘pure’ when independent of genotypes [27]. The most commonly studied type of epigenetic variation at the population level is DNA methylation, a biochemical modification of the DNA sequence adding a methyl group, generally to a cytosine within a CpG dinucleotides [28]. Level of DNA methylation in regulatory regions can influence the level of gene expression, with generally a negative correlation between DNA methylation level and gene expression [28], although opposite pattern has been documented with an increase methylation in core genes [29]. Consequently, DNA methylation level associated with a change in gene expression of the gene underpinning an adaptive phenotypic trait under selection might contribute to generate an adaptive phenotypic response [30].

Here, we compared two sister species complexes of whitefish (Lake whitefish: *Coregonus clupeaformis* and European whitefish: *C. lavaretus*), both comprising sympatric benthic and limnetic specialists [31,32]. The Lake whitefish and the European whitefish (hereafter, lineages) evolved separately on both continents since they became geographically isolated ∼500,000 years ago [33-35]. The limnetic-benthic species complex evolved independently on both continents in two separate glacial lineages during the last Pleistocene [33,36-39], and on both continents, they result from a post-glacial secondary contact which then colonized independent post-glacial lakes [39,40]. The limnetic species colonized the free limnetic ecological niche through an adaptive divergence with an associated phenotypic evolution that translated into slower growth, slender body, higher metabolic rate and more active swimming behaviour [41-46], that relied on convergent genetic and transcriptomic bases [8,47-49], and are reproductively isolated from the sympatric benthic species [50]. As such, this whitefish species complex offers a valuable model to study the molecular bases associated with convergent phenotypic differentiation.

The main goal of this study was to document the relative role of genomic vs. epigenomic variation in shaping patterns of phenotypic variation, particularly at the gene expression level during independent ecological speciation events. We first tested for convergence in DNA methylation differentiation between limnetic and benthic whitefish from both North America and Europe to test for the occurrence of non-random epigenetic associated limnetic-benthic diversification. Next, we combined transcriptomes and whole epigenomes resequencing data to assess the effect of DNA methylation on phenotypes (here, gene expression) associated to the limnetic-benthic diversification. Then, we identified eQTLs allowing to compare the relative effect of both genomic and epigenomic variation on patterns of convergent gene expression differentiation between limnetic and benthic species from both continents. Finally, we estimated the proportion of i) ‘pure’ genetic effects, ii) putatively ‘obligatory’ and ‘facilitated’ epigenetic effects (that we will thereafter call ‘genomic-epigenomic’ interactive effects), and iii) ‘pure’ epigenetic effects on phenotypic diversification during speciation.

## RESULTS

### Convergent differential gene expression between limnetic and benthic species

We sampled two lakes in North America (USA: Cliff Lake and Indian Lake) and two lakes in Europe (Norway: Langfjordvatn Lake and Switzerland: Zurich Lake), presenting a sympatric limnetic-benthic species pairs (Fig. 1). Liver tissue RNAseq generated a total of 1.15×10^9^ 100bp raw single-end reads from 48 individuals (six individuals per species for a given lake; Table S1). Filtered libraries (1.13×10^9^ reads) were aligned to the reference transcriptome to quantify the amount of reads per transcript per individuals. We quantified differentially expressed genes (DEG) between conditions (*i*.*e*., limnetic and benthic species) using a generalized linear model, including lake, continent and species information as covariates. We found 189 convergent DEGs (*Bonferroni* < 0.1) between all limnetic vs. all benthic whitefish. These convergent genes showed a higher proportion of down-than up-regulated genes in limnetic species relative to the benthic species (68 vs. 121; χ^2^ = 14.862, df = 1, *P*=0.0001). Among the overexpressed genes, four of them have been annotated as transposable elements (TEs, Table S2).

**Fig. 1.**
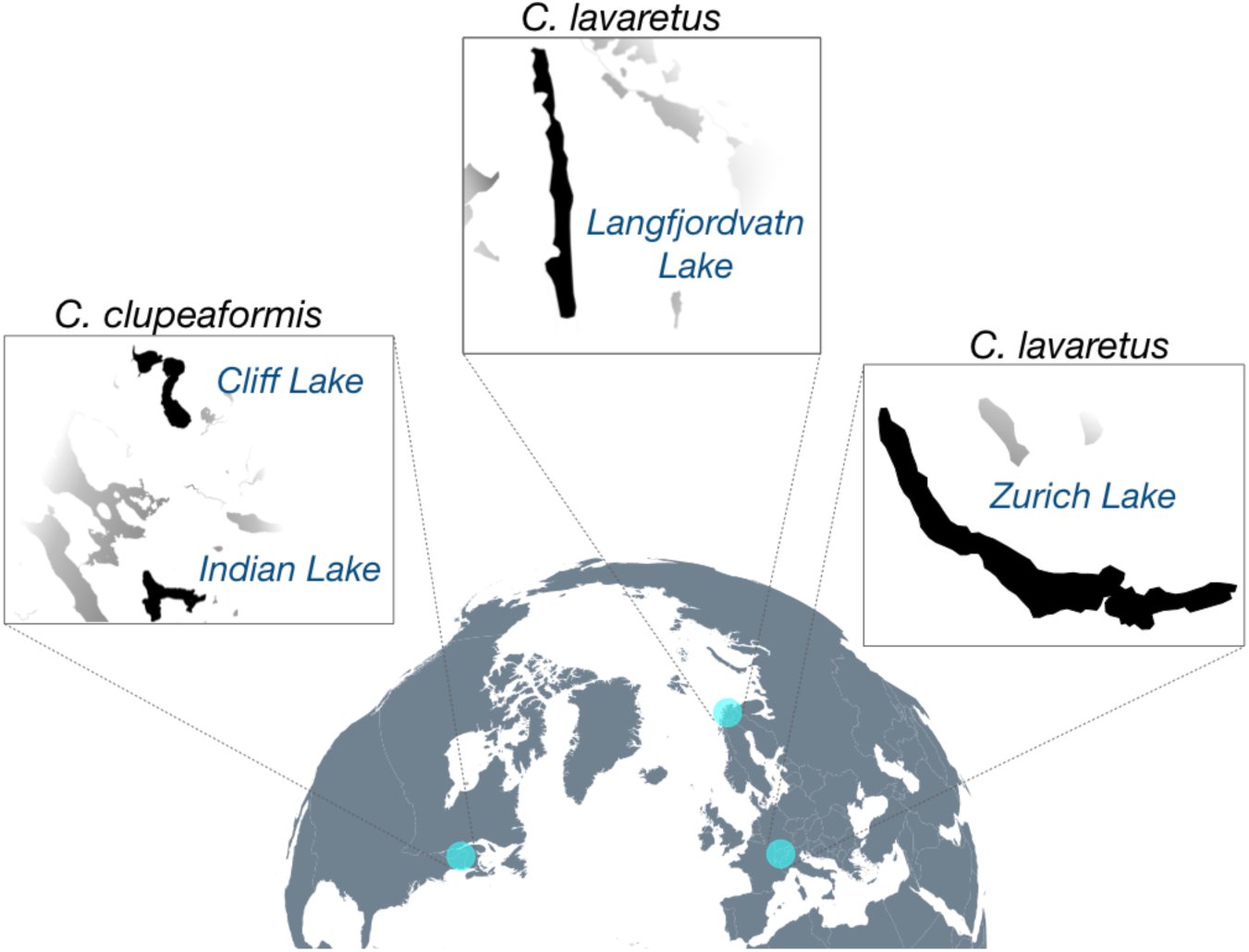
Geographic location studied lakes harbouring sympatric limnetic and benthic species. Two lakes were sampled in North America: Cliff Lake and Indian Lake in the region (blue circles) of Maine, and two lakes were sampled in Europe: Langfjordvatn Lake in Norway and Zurich Lake in Switzerland.

Gene ontology enrichment analysis revealed an overrepresentation of modules associated with metabolic processes (GO:0006807, nitrogen compound metabolic process; GO:0044237, cellular metabolic process; GO:0044238, primary metabolic process; GO:0071704, organic substance metabolic process; GO:0008152, metabolic process), immune system response and antioxidant activity (GO:0016209, antioxidant activity; GO:0003823, antigen binding) and methylation (GO:0032259, methylation) in limnetic species (Table S3).

### Convergent differential methylation between limnetic and benthic species

A total of 10.9×10^9^ 150bp paired-end reads were generated from 64 individuals for the WGBS, resulting in an average coverage of 13.6X (s.d. 1.6X) per individual (Table S4). Those 64 individuals were the same as those previously described for transcriptomic analyses, with two individuals per species added for a given lake. After filtering for C-T genetic polymorphism and CpGs corresponding to CGs context, the average number of CpGs among all populations was 949,828 (s.d. 114,690). The number of differentially methylated loci (DML; significant DNA methylation differentiation at the same position) between limnetic and benthic species was 18,266 DMLs (s.d. 3,829) on average and the number of differentially methylated regions (DMR; cluster of at least three DMLs in a minimum of 50bp) between species was 619 DMRs (s.d. 93) on average (Table S5).

We then tested for the occurrence of convergent differential methylation between all limnetic and all benthic fish across both continents. Generalized linear models identified 149 significant DMRs between all populations of the limnetic species and all populations of the benthic species across both continents that were shared among all populations. From the 149 DMRs, approximately twice as many DMRs were hyper-methylated in the limnetic species relative to the benthic species (92 hyper-methylated *vs*. 57 hypo-methylated; χ^2^ = 8.22, df = 1, *P* = 0.004), which suggests a non-random epigenetic pattern underlying convergent phenotypic differentiation between limnetic and benthic whitefish across both continents.

The biological functions associated with hyper-methylated DMRs were very distinct between limnetic and benthic species across both continents. Thus, the analysis of gene ontology enrichment of the 92 hyper-methylated DMRs in the limnetic species showed an overrepresentation of biological processes linked to growth and developmental functions (GO:0048856, anatomical structure development; GO:0032502, developmental process; GO:0044699, single-organism process; GO:0009987, cellular process), while the 57 hyper-methylated DMRs in the benthic species showed enrichment in genes associated to variable rate of cell cycle process (GO:0022402 BP, cell cycle process) (Table S6).

### DMRs are associated with gene expression

The 57 hypo-methylated DMRs in the limnetic species were mainly found in overexpressed genes relative to the benthic species (*i*.*e*., positive log2 Fold change; Fig. 2A, Fig. S1) and conversely, hyper-methylated genes in limnetic species showed a lower gene expression relative to the benthic species (Fig. 2A). This difference in expression between hyper-*vs*. hypo-methylated DMRs genes was highly significant (W = 4577, *P* < 0.001). Finally, we found a strong positive correlation (r = 0.91, df = 2, *P* = 0.08) between the proportion of repressed genes and the significance level of DMRs (Fig. S2).

**Fig. 2.**
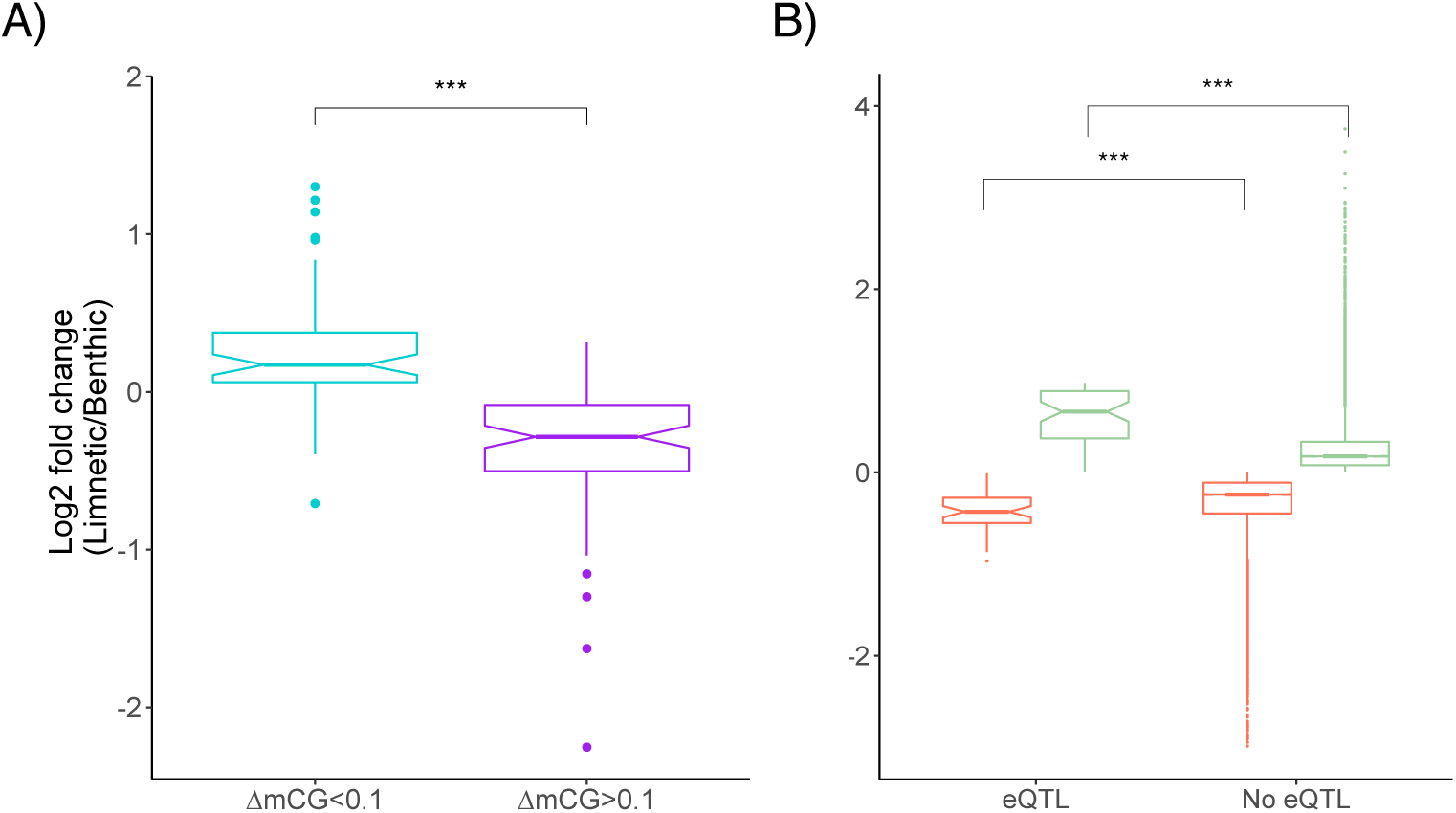
Differential gene expression between limnetic and benthic whitefish induced by DNA methylation and cis-eQTL. A) Boxplot showing the level of gene expression differentiation (Log2 fold change) as a function of the difference in methylation level between the limnetic and benthic species (ΔmCG Limnetic-Benthic) for genes associated with a convergent DMR. The light-blue box corresponds to genes with hypo-methylated DMRs in limnetic species, whereas the purple box corresponds to genes with hyper-methylated DMRs in the limnetic species. Both categories are respectively associated with an overexpression (Log2 fold change > 0) and a repression (Log2 fold change < 0) of gene expression and showed a significant difference in the level of gene expression (*P* < 0.001). B) Boxplot showing the level of gene expression differentiation (Log2 fold change) as a function of genes for which we found a presence (eQTL) or absence (No eQTL) of convergent *cis*-eQTL. The orange and light-green boxes correspond to repressed and overexpressed genes, respectively, in either gene category with *cis*-eQTL or without *cis*-eQTL. Overexpressed genes in limnetic species (Log2 fold change > 0) are more differentiated when affected by a *cis*-eQTL (*P* < 0.001), as well as repressed genes (Log2 fold change < 0) (*P* < 0.001).

### eQTL effect on differential gene expression between species

We then associated SNP variation for a given gene and its level of expression while correcting for population structure using a generalized linear model (glm). We found 108 significant convergent *cis*-eQTLs at a false discovery rate (FDR) of 0.01 among all limnetic and all benthic whitefish (Fig. S3). From the 108 convergent transcripts harbouring a *cis*-eQTL, 17 of them were identified as DEGs (hypergeometric test, *P* < 0.001). Moreover, none of the genes harbouring a *cis*-eQTL had a DMR, which therefore represents a measure of genetic effects, which could be compared to epigenetic effects. Thus, we first contrasted the variance in gene expression between limnetic and benthic species in genes for which a *cis*-eQTL was associated and for genes without *cis*-eQTL. We found that for repressed genes, those without *cis*-eQTL showed a more pronounced variance in gene expression compared to those with a *cis*-eQTL (F = 0.3723, num df = 49, *P* <0.001). However, no significant difference in the gene expression variance was observed in overexpressed genes (F = 0.7936, num df = 57, *P* = 0.26), in the limnetic species relative to the benthic species. We also observed a significant difference between genes with and without *cis*-eQTL, either for genes that were repressed (*i*.*e*., negative Log2 fold change; F = 4.1309, num df = 88, *P* < 0.001) or overexpressed (F = 10.434, num df = 56, *P* < 0.001; Fig. 2B) in limnetic relative to benthic species. In addition, the difference in levels of gene expression between limnetic and benthic species was more pronounced for genes with cis-eQTL compared to genes without cis-eQTL, both in repressed (Wilcoxon test, W = 323420, *P* < 0.001), and overexpressed (Wilcoxon test, W = 647930, *P* < 0.001) genes in limnetic whitefish. Moreover, genes with cis-eQTL (genetic effect) showed a more pronounced difference in gene expression between limnetic and benthic species than those with DMR (epigenetic effect), both in repressed (Wilcoxon test, W = 2797, *P* = 0.012), and overexpressed (Wilcoxon test, W = 677, *P* < 0.001) genes in the limnetic whitefish (Fig. S4).

### Genomic and epigenomic effects on gene expression

Redundancy analyses were produced to partition the variance in gene expression that was explained by genomic and epigenomic components. When we considered the entire dataset, genomic and epigenomic explained together 53.1% of the variance in gene expression (*P* < 0.001; Fig. 3A). This quantitative assessment of the contribution of genomic and epigenomic was inferred through PCAs for gene expression, genomic and epigenomic, that showed similar patterns by separating continents and lakes (Fig. 3B, C and D, respectively; Table S7). Genomic and epigenomic components separately explained 50.8% (*P* < 0.001) and 49.0% (*P* < 0.001) of the variance in gene expression, respectively (Fig. 3A). However, when controlling for epigenomic, 4.1% (*P* < 0.05) of gene expression variation was explained by ‘pure’ genomic effects, and when controlling for genomic, only 2.3% (*P* < 0.1) was explained by ‘pure’ epigenomic effects (Fig. 3A), and 46.7% of the variance in gene expression was thus explained by both components that we coin as putatively genomic-epigenomic interactive effect (Fig. 3A). Therefore, the model did not explain 46.9% of the variance in gene expression (Fig. 3A), which could be attributed to the lack of the entire promotor region in most of the sequenced genes, to other regulatory machineries (*e*.*g*., *trans*-regulation) and/or epigenetic mechanisms other than DNA methylation. The partitioning of gene expression variance by genomic and epigenomic was reproduced within each continent. Similar results were found, except for the ‘pure’ epigenomic effect, which was not significant on either continent (*P* > 0.1; Fig. 3E and I for America and Europe, respectively; Table S7).

**Fig 3.**
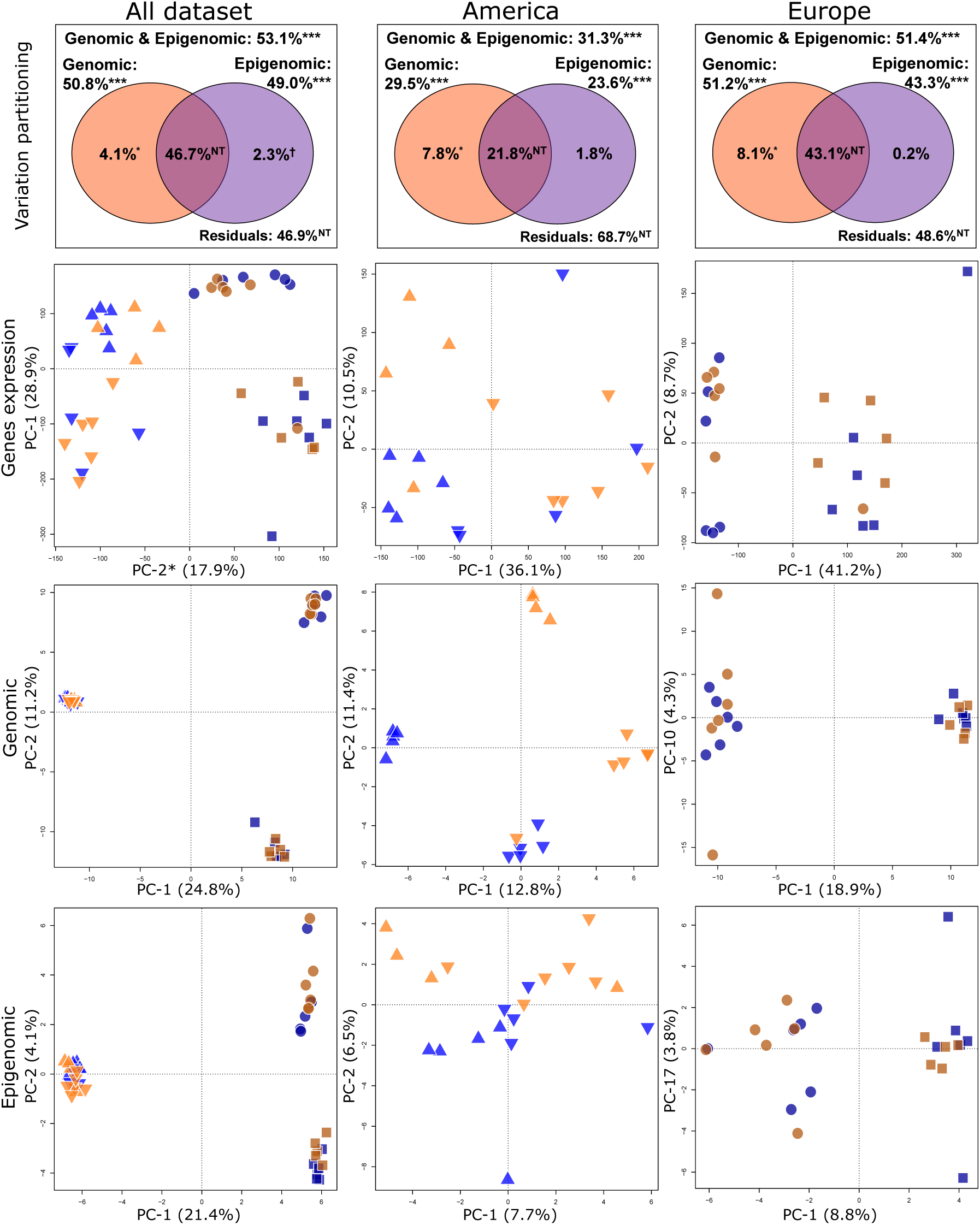
Partitioning of the variance in gene expression and epigenomic factors. The figure summarizes the variance partitioning of orthologous genes expression associated to genomic (*i*.*e*., trans-specific polymorphism) and epigenomic (*i*.*e*., trans-specific CpGs) data, on panels of the first row, for limnetic-benthic species comparisons across both continents (All dataset, first column), and for limnetic-benthic species comparisons in *C. clupeaformis* (North Amerixa, second column) and *C. lavaretus* (Europe, third column). Each panel of the variation partitioning decomposes the variance in gene expression measured in each population of the limnetic and benthic species, at both the continental and lake scale. The total amount of variance explained by the data corresponds to the ‘Genomic & Epigenomic’ category, while the remaining part is associated to the statistically non-testable (NT) ‘Residuals’. Then, the explained proportion of variance in gene expression is decomposed by genomic factor alone (‘Genomic’) and the epigenomic factor alone (‘Epigenomic’). The proportion of variance associated to ‘Genomic’ (darkgreen), ‘Epigenomic’ (purple) and their intersection (dark purple, middle) are decomposed in the Venn diagram. Other panels illustrate the variance between individuals within populations of both species resolved using PCAs, for the three biological levels (‘Gene expression’, second row; ‘Genomic’, third row and ‘Epigenomic’, fourth row). For each PCA, the plotted axes tied in to axes identified by backward selection, and each symbol corresponds to an individual fish. Blue and orange symbols represent limnetic and benthic species, respectively. Lighter tones for both colors are for *C. clupeaformis* (North Amerixa) individuals and darker tones for *C. lavaretus* (Europe) individuals, whereas different shapes represent Cliff (triangles), Indian (inverted-triangles), Langfjordvatn (circles) and Zurich (squares) lakes. *Note that on the gene expression PCA for the ‘All dataset’, we inverted the axes (PC-1 and PC-2) during the projection for readability and comparison of patterns with other PCAs.

## DISCUSSION

Our study focus on two sister lineages of whitefish that have diverged and evolved independently for about ∼500,000 years [34] and have postglacially colonised cold freshwater lakes of Eurasia and North America, approximately 12,000 years ago, since then [33,36,51,52]. In both lineages, diverging limnetic and benthic sympatric species pairs are associated with their respective trophic niches [53-55]. The repeated sympatric convergent phenotypic diversification occurred in several lakes where whitefish have undergone a secondary contact between diverged glacial lineages (*i*.*e*., sub-lineages that were formed in allopatry on each continent during the last glaciation event) [39,40]. In this study, whole transcriptome and epigenome data offered the opportunity to analyze both the level of DNA methylation and gene sequence divergence and combine these to the analysis of differential gene expression on representative sympatric pairs of limnetic-benthic whitefish species from two continents. This allowed addressing fundamental questions pertaining to the relative importance of different molecular mechanisms associated with the phenotypic differentiation in a context of repeated ecological speciation.

### Functional effects of convergent DEGs

The identified convergent DEGs indicate that divergence between replicated species pairs act on a partially shared set of genes. First, we found consistent patterns of convergent DEGs annotated as TEs between limnetic and benthic species, supporting a role for such genomic component in the gene expression variation between species [56], or the existence of annotation errors due to the widespread presence of TEs in salmonids genomes. Second, convergent DEGs between species across the entire system were significantly enriched in metabolic processes and immune functions. This corroborates previous studies on North American whitefish species pairs in which the authors proposed that the differences in both metabolic and immune functions between limnetic and benthic whitefish actually reflect life history trade-offs. These trade-offs may involve a more active swimming in order to avoid predation and an increased foraging efficiency in dwarf whitefish, possibly counterbalanced by an increased probability of parasite infection and consequently, increased mortality risk and increased energetic costs translating into slower growth [8,41,46,48,54,57]. DEGs were also enriched in genes associated with methylation regulation, suggesting a genetic role of DNA methylation. More precisely, the enrichment for methylation regulation was associated to the process of rRNA 2’-O-methylation (rRNA2’-O-me), a highly complex and specific posttranscriptional modification present in functionally important domains of the ribosome (Krogh *et al*. 2016). Such mechanism within the ribosome is particularly important in the regulation of the gene expression [58]. Moreover, some functional domains of ribosomes have been shown to be targeted by the rRNA2’-O-me, inducing a modulation of the translation of mRNAs through plastic response [59]. Thus, such mechanisms suggest an indirect gene expression regulation through DNA methylation.

### Origin of convergent DMRs

A salient result of our study is the non-random observation of DMRs between limnetic and benthic whitefish across the system, and more specifically the occurrence of 149 shared DMRs across continents between all limnetic *vs*. all benthic whitefish (convergent DMRs). The identification of convergent DMRs between all limnetic and benthic whitefish, from independent populations distributed across two continents, suggests that epigenetic differentiation is likely to be mostly influenced by environmental pressures associated with the use of different ecological niches by limnetic and benthic whitefish [60,61]. Convergent DMRs could reflect a direct response to local ecological conditions [62,63] or be directly associated to the modulation of the gene expression by genetically induced methylation [27]. The later suggest that genetic [64] or the interplay between genetic and environment [64] could have led to such methylation convergence between replicated species pairs across continents. Here, the weak/no proportion of variation in gene expression explained by ‘pure’ epigenetic component (see below for a further discussion) support this hypothesis. Admittedly however, rigorously testing this hypothesis regarding the genetic origin of DMRs would require performing experimental crosses between species in common garden experiments. We recently performed such a controlled experiment on North American limnetic and benthic whitefish which revealed limited phenotypic plasticity when reared in contrasting environments [41,46]. Similarly, common garden experiments on the same system revealed that differential gene expression [48], and high methylation differentiation that was strongly associated with the presence of transposable elements (TEs) was maintained when limnetic and benthic whitefish were reared in the same environmental conditions [52]. DNA methylation silence TEs activity and TEs activity have been shown to affect nearby gene expression, suggesting that their interplays can contribute to phenotypic differentiation [56]. In our empirical analysis, despite the lack of significant DMR in the convergent differentially expressed TEs between species across the system, those TEs harboured significant DMLs that were hypo-methylated in the limnetic species. This suggests a putative role of DMLs in the local transcriptomic regulation. Interestingly, genes exhibiting persistent trans-generational DNA methylation patterns often contain a TE insertion nearby [65]. Together, this supports the hypothesis that convergent DMRs observed in this study may have a genetic basis rather than merely reflecting the influence of local environment.

### Functional effect of convergent DMRs

The gene ontology analysis of genes with convergent DMRs revealed an enrichment of gene modules involved in growth and developmental processes, and more so for hyper-methylated DMRs in the limnetic species, but also genes module regulating cells cycle process for hypo-methylated DMRs in the limnetic species. These results suggest a repression of expression of genes related to growth and developmental process in the limnetic species while genes associated with a higher metabolism and shorter cell cycle life are overexpressed in the limnetic species relatively to the benthic species. These observations at the epigenetic level corroborate those of previous studies at the transcriptomic and physiological levels performed on these populations and further supports the hypothesis of life history trade-offs between survival (limnetic) and growth (benthic) functions revealed in our previous transcriptomic studies [8,48,66-68]. On the other hand, none of the convergent DMRs belong to a DEG, suggesting that methylation differentiation is not a major mechanism involved in the highest differential of gene expression associated to the divergence of limnetic and benthic species. Similar observations have been made in the stiff brome (*Brachypodium distachyon*), where CpGs, DMRs and gene expression associations were not systematically related across tissues comparisons [69], as well as in hepatocellular carcinoma cells where differentially expressed genes were not correlated to higher DNA methylation differences [70]. Here, the transcript-associated convergent DMRs were involved in biological functions associated with differential phenotypic and ecological differences between limnetic and benthic whitefish on both continents.

### Effects of DNA methylation on gene expression

We then investigated the possible association between differential patterns of gene expression with the extant of methylation for those genes that showed convergent DMRs between all limnetic and all benthic whitefish. Most of the convergent hypo-methylated DMRs identified in the limnetic species were associated to overexpressed genes, compared to the benthic species, while convergent hyper-methylated DMRs in the limnetic species were associated with a repression of the gene expression. These results support the correlation between DNA methylation level and the level of transcriptional activity of linked genes, where hyper-methylation is typically associated with gene expression repression [28,71]. In addition, the most significant DMRs were directly associated with a higher proportion of genes with a repressed transcriptional activity. Moreover, patterns of methylation arose from the density of DMLs within DMRs but mainly, from the difference in the proportion of methylated sites cumulated within DMRs (Fig. S2, S5) and the amount of methylation differences along the DMR between species, as previously observed [72].

### Effect of genetic vs. epigenetic variation on gene expression

Another salient observation supporting the prevailing role of genetic over epigenetic variation is that differences in gene expression between limnetic and benthic whitefish was more important for genes for which we identified a *cis*-eQTL than the rest of the genes (*i*.*e*., without genetic and with or without epigenetic effects). It is also important to point out that none of the genes harbouring a *cis*-eQTL were associated with a DMR. This pattern emerged for overexpressed and repressed genes in the limnetic species relative to the benthic species. Also, both gene categories with *cis*-eQTL showed less gene expression variance than those without *cis*-eQTL. This observation also suggests that selective pressures act on standing genetic variation (*i*.*e*., shared variants across the system) and that favoured alleles could increase in frequency, as previously described in the whitefish system [8]. Similar patterns were also observed in other taxonomic groups. For example, *cis*-variation in a gene promoter can alter the binding site of a regulatory protein and induce newly derived phenotype. This was observed in the morphological evolution of two *Drosophila* species [73], in the three-spine stickleback where genetic variants in the EDA gene are associated with the presence/absence of armour plates [74] or *cis*-acting regulation between divergent marine-freshwater ecotypes [17], as well as in corals under changing climatic conditions where the frequency of favoured alleles co-vary with gene expression level [75]. Moreover, genes affected by *cis*-eQTL showed higher gene expression differentiation between limnetic and benthic species than genes with convergent DMRs between limnetic and benthic whitefish, and more so for overexpressed genes in the limnetic species relative to the benthic whitefish. Furthermore, despite the effect of methylation on their expression, genes associated with convergent DMRs were not associated with most DEGs whereas those associated with *cis*-genetic variants were (15%) [8,47]. These qualitative observations suggest that variation in gene expression can result from direct changes in the level of DNA methylation but this effect appears less important than genetic *cis*-acting variants.

This hypothesis was supported by the multivariate statistical framework developed to disentangle the relative role of genomic and epigenomic variation on patterns of gene expression. Indeed, combined data set (*i*.*e*., genomic and epigenomic) explained 53.1% of the variation in gene expression between species across continents, with 4.1% attributable to ‘pure’ genetic, 2.3% to ‘pure’ epigenomic and 46.7% to putatively genomic-epigenomic effect. Thus, the observed variance in gene expression variation was weakly explained by ‘pure’ epigenomic effects (*i*.*e*. 0.05 < *P* < 0.1). Similar qualitative and quantitative observations of a lower contribution of DNA methylation compared to genetic variation were made in previous studies. For example, Ahsan *et al*. investigated the relative contribution of DNA methylation and genetic variants on biomarkers of human diseases, and found that for 36% (44/121) of the studied proteins the abundance variation relied on DNA methylation bases while 52% (23/44) of these were associated to genetic variants (*i*.*e*., interaction between DNA methylation and genetic) [76]. Moreover, studies focusing on the bases of adaptive traits in *Arabidopsis thaliana* and combining both DNA methylation and genomic, highlighted that DNA methylation variation was less likely to contribute to the adaptive traits than standing genetic variation [64,77]. Overall, these quantitative results confirmed that variation in gene expression relied mainly on hard-coded genetic bases, and that most of the epigenomic associated to gene expression variance are in direct interaction with genomic effects. This suggests that methylation patterns observed among generations could be attributable to the inheritance of some associated genetic variation.

### Limitations of this study

Our experimental design, while relevant to address questions pertaining to the study of convergence due to repeated phenotypic diversification between limnetic and benthic species, relied on individuals sampled from the wild. Such characteristics could involve increased inter-individual variation of the DNA methylation level thus reducing the amount of CpGs potentially associated to DMLs or DMRs detected in our statistical analyses.

Moreover, our work focuses on liver tissue DNA methylation levels and comparisons between individuals from different populations of two species. The use of this tissue allows direct comparisons with previous transcriptomic studies, comparing limnetic and benthic species [8,48,78]. This tissue is particularly interesting because most of the energetic and metabolic genes are expressed, and are relevant to study metabolisms (*e*.*g*., development, energy) associated the adaptive phenotype diversification of species to different ecological niches [79]. While it could have been ideal to realize this study on several tissues, such design was not realistic in terms of costs. Therefore, we chose to work on liver tissue because of its homogeneous tissue characteristics, allowing a reduced variation in DNA methylation between cell lines, and because more genes are expressed in this organ than most other tissues or organs studied in fishes [66] and in human [80,81], Then, despite the increasing number of studies aiming to identify the origin of epigenomic variant and their relative effect, difficulties remained to identify pure epigenetic variation as response to change in environmental conditions, as observed in some systems [82,83]. Furthermore, the mechanism of heritability of epigenetic variations has not been documented in whitefish, and no mechanism of environmentally induced variation transmitted to offspring has been highlighted properly in the wild [83,84], contrary to theoretical work [30], notably due to a lack of knowledge in mechanisms associated with heritability [26,85]. Thus, our knowledge on the relative effect of pure epigenetic variation on the evolution of wild populations is limited, although obligatory epigenetics affecting the fitness of a trait could influence the evolution of populations [27,86].

Finally, here we used a transcriptome as genomic reference. Then our study focused mainly on coding regions and partial flanking regions. In our statistical framework, 46.9% of gene expression was not associated to methylation and/or genomic variation. This observation suggests that other mechanisms associated to gene expression regulation such as *trans*-acting genetic and epigenetic variants that likely act directly on enhancers and/or transcription factors, as well as other epigenetic process (*e*.*g*., histone structure and small RNAs) and other genomic variation (e.g., structural genomic variants), that are still to be investigate to better understand the role of epigenetic in speciation [87,88]. Future works on the whitefish system should improve in resolution by the use of the newly assembled reference genome for the *C. clupeaformis* species (unpubl. Data). Then, the different gene regions (*i*.*e*., UTRs, promotor, introns and exons) should be analysed separately in order to quantify the level of methylation along genes and focus on their relative effects on gene expression levels.

### Concluding remarks

In this study, we have reported a qualitative and quantitative assessment of the role of genomic and epigenomic factors and their interactive effect on gene expression variation, and identified the origins of the methylation variation in a non-model species. We observed that gene expression differentiation between diverging species pairs was mostly caused by *cis*-genetic variants or genomic-epigenomic interactive effect. Interestingly, similar observations emerged from previous studies on model species. For instance, several studies focused on the inheritance of CpG methylation sites among human family cohorts and identified that most of the epigenomic variation were inherited in Mendelian proportions, and that only 3% of CpGs were not associated to a genetic basis [84]. This supports the idea that the majority of epigenomic variation have a genetic basis, which corresponds to the definition of ‘obligatory’ or ‘facilitated’ methylation [27]. Theory predicts that ‘pure’ epigenetic variation could allow an independent selection from the genetic matrix on newly induced phenotypes [30]. Yet, our study, along with previous empirical ones mainly point to a marginal contribution (if any) of ‘pure’ epigenetic effects compared to the contribution of genetic and genetic-associated epigenomic variants. For example, testing for environmentally-induced methylation in *Arabidopsis thaliana* coupled with and extensive genome wide association study (GWAS) revealed that most of the methylation variation was associated to changes in locally adapted allele frequencies, thus ruling out ‘pure’ epigenetic contribution [64]. Moreover, comparisons between human populations revealed methylation modifications with a strong genetic basis but a weak contribution of epigenetic variation on gene expression [89]. Consequently, we believe that more caution should be taken when interpreting epigenetic (*i*.*e*., methylation) patterns to infer evidence for local adaptation when the origin of such epigenetic variation is not initially defined or controlled for the genomic component.

## MATERIALS AND METHODS

### Sample preparation, transcriptomes and whole genomes bisulphite sequencing

We sampled whitefish from two lakes in North Amerixa (Cliff Lake and Indian Lake) and two lakes in Europe from Norway (Langfjordvatn Lake) and Switzerland (Zurich Lake), each comprising sympatric limnetic-benthic species pairs (Fig. 1). Six individuals per species per lake were sampled, for a total of 48 individuals. DNA and RNA were extracted from liver tissue of each sample, stored in -80°C and in RNA later (All samples from Europe were stored in RNA later). Whole genomic DNA was isolated using the DNeasy Tissue Kit (Qiagen, Valencia, CA), DNA integrity was checked on an agarose gel (1%), and quantified with optic density measure (NanoDrop™2000, Thermo Scientific, Waltham, MA) before performing quantitative-PCR. Individual libraries were built at the McGill University and Genome Quebec Innovation Centre (Montreal, Canada), for Illumina paired-end 150bp whole-genome bi-sulfite sequencing (WGBS) on the Illumina HiSeqX. The libraries for the WGBS were combined randomly on 16 lanes (two flow-cells, four individuals pooled per lane for initially 64 libraries). From six of these eight individuals per population, we extracted the total RNA from liver tissue as detailed in [8]. Briefly, we chose the liver tissue for i) its homogeneous tissue characteristics and because more genes are expressed in this organ than most other tissues or organs studied in fishes [66] and in human [80,81], ii) its multiple biological functions such as growth regulation in Salmonids [90], but also in energy metabolism, iron homeostasis, lipid metabolism and detoxification which show heritable divergence in limnetic and benthic species [48]. Before any handling, each sample for RNAseq analysis was assigned to a random order. RNA was extracted using the RNeasy Mini Kit following the manufacturer’s instructions (Qiagen, Hilden, Germany). RNA quantity and quality were assessed using the NanoDrop™2000 (Thermo Scientific, Waltham, MA), and the 2100 Bioanalyser (RIN > 8.0) (Agilent, Santa Clara, CA, USA). Single read sequencing (100bp) was performed on the Illumina HiSeq 2000 platform for the 48 libraries (initially 72 libraries including other populations used in [8]), at the McGill University and Genome Quebec Innovation Centre (Montreal, Canada).

### Gene expression analysis

Gene expression analysis was realized as in a previous study [8]. However, we realized an original gene expression comparison based on a log-likelihood ratio test, as detailed below. Raw sequences were cleaned from adaptor and tag sequences, and trimmed using *Trimmomatic* v0.36 [91]. Individual reads were mapped to the reference transcriptome using Bowtie2 v2.1.0 [92]. *eXpress* v1.5.1 [93] was used to estimate individual reads counts from BAM files. We projected the raw estimated counts in order to control for (if any) pattern of batch effect (absence of such patterns, Fig 3b). Then, the analysis of differential expression was performed with *DESeq2* v1.14.1 [94]. Counts matrix was normalized using size factors and was log2-transformed. Considering the hierarchical structure of the studied system, we used a log-likelihood ratio test (LRT) approach in order to test for the effect of species (benthic species determined as reference), by building a full general linear model allowing species comparisons while integrating lakes and continents effect as covariates, and controlling for structuration with a reduced model. A significant threshold of a FDR < 0.1 was applied to determine differentially expressed genes (DEGs). Additional models testing for comparisons between continents and lakes only were tested in order to control for DEGs associated with environmental effects (*e*.*g*., lakes or continent) and we discarded transcripts in interaction/intersection with any environmental effect.

### SNP calling

Raw data from the 48 transcriptomes were cleaned and trimmed using cutadapt (v1.10) [95]. Cleaned reads were aligned to the indexed reference transcriptome [8] with *Bowtie2* v2.1.0 [92]. SAM files obtained were converted to BAM files, sorted using *Samtools* v1.3 [96] and cleared from duplicates with the *Picard-tools* v1.119 program (http://broadinstitute.github.io/picard/). Reads alignment information to the reference transcriptome was used for genotyping SNPs with a minimum of quality of alignment of four and a minimum number of five reads to call SNPs with *Freebayes* v0.9.10-3-g47a713e [97]. *Vcffilter* program from *vcflib* [98] was used in order to keep variable sites with a minimum coverage of three reads per individual, bi-allelic SNPs with a phred scaled quality score above 30, a genotype quality with a phred score higher than 20. Then, we filtered the resulting VCF file using *VCFtools* [99], in order to remove miscalled and low quality SNPs for subsequent population genomics analyses. For each of the eight populations, we kept loci with less than 10% of missing genotypes and filtered for local MAF (0.01 per population). Finally, we merged the VCF files from all eight populations, resulting in a unique VCF file containing 161,675 SNPs passing all the filters, in order to correct WGBS data (see below), and a VCF file of 9,093 SNPs shared among all populations across continents (trans-species polymorphism) and with no missing data for subsequent analysis.

### Methylation calling and differential methylation analyses

Raw sequencing reads were trimmed for quality (≥25), error rate (threshold of 0.15) and adaptor sequence using *trim_galore* v0.4.5 (http://www.bioinformatics.babraham.ac.uk/projects/trim_galore/). Trimmed sequences were aligned to the reference transcriptome using *BSseeker2* v2.1.5 [100] with *Bowtie2* v2.1.0 [92] in the end-to-end alignment mode. We cleaned the BAM files by removing duplicates with the *Picard-tools* v1.119 program (http://broadinstitute.github.io/picard/), before determining methylation levels for each site by using the *BSseeker2* methylation call step. The raw methylation file was filtered by removing C-T DNA polymorphism identified from ‘SNP calling’ step, in order to avoid ‘false’ methylation variation at those positions [101]. We used the *CGmapTools* suite v0.1.1 [102] to extract only CpGs sites determined as CG context (avoiding CHH and CHG contexts), and with a minimum of 10X coverage and a maximum of 100X in order to avoid noise from repetitive elements and paralog genes. Generalized linear model, with the hierarchical population structure as covariate, was used to identify differentially methylated loci (DMLs) and differentially methylated regions (DMRs) with the *DSS* R package [103], using a smoothing strategy on 500bp. We also controlled for interaction between terms allowing direct comparisons between species. Then, we compared directly all limnetic individuals to all benthic individuals across continents. DMLs were defined when showing at least 20% of difference between species and a significant threshold of *P* < 0.05. DMRs were retained when at least five CpGs occurred in a minimum sequence of 50bp, when CpGs showed a minimum of 10% of methylation difference between species and a significant threshold of *P* < 0.05. DMRs distant by 50bp or less were merged together to be defined as the same DMR.

### Gene ontology analyses

Gene ontology (GO) enrichment analyses were performed on gene expression and methylation with *GOATOOLS* [104]. We tested significant DEGs and DMRs using Fisher’s exact tests and GO enrichment were associated with FDR < 0.05 (Benjamini-Hochberg correction) and we kept GO categories represented by at least three genes.

### Effects of genetic and DNA methylation on gene

Gene expression level was associated to sequence polymorphism to identify eQTLs. We used the R package *MatrixEQTL* v2.1.1 [105] to perform association mapping for local eQTL affecting the expression level of the transcripts to which they were directly physically linked (*cis*-eQTL). From the 9,093 shared SNPs among all eight populations, we retained loci showing polymorphism across continents (*i*.*e*., existence of the three genotypes for a given position among all studied populations), which corresponded to 5,424 SNPs. eQTLs were identified through linear models in order to identify differential expression between species, while considering the different hierarchical levels as covariates (*i*.*e*., species, lakes and continents). A false-discovery-rate correction was applied and significance of identified *cis*-eQTL was accepted with a FDR<0.01.

Then, we tested for association between levels of gene expression and variation of genes affected by a convergent DMR across continents. Considering convergent DMRs, only one DMR per gene was retained, keeping the DMR with the higher number of CpGs between species. We then directly associated the level of differential methylation between limnetic and benthic species to the difference in expression (Log2 Fold Change) for the gene to which the DMR is associated to. Indeed, it has been showed that the evaluation of the relationship between methylation level and gene expression, considering the log fold change is a more relevant signal of change in gene expression than the absolute differences [106]. This approach allowed defining genes with and without DMRs and *cis*-eQTL.

### Variance of gene expression explained by genomic and epigenomic

We produced three redundancy analyses (RDAs) to estimate the percentage of gene expression variation explained by i) genomic variation alone, ii) epigenomic variation and iii) both molecular mechanisms. Then, two partial RDAs (pRDAs) were applied for quantifying the proportion of variance in gene expression explained by i) genomic variation, when controlling for epigenomic variation (thereafter call ‘pure’ genomic), and ii) *vice versa* (thereafter call ‘pure’ epigenomic). Then, we defined the epigenomic component relying on genomic bases (namely ‘obligatory’ and ‘facilitated’ epigenetic) as genomic-epigenomic interactive effect. Estimation of gene expression variance explained by genomic-epigenomic interactive effect was computed with the function ‘varpart’ available in the *Vegan* R package [107]. RDAs and pRDAs were performed using the function ‘*rda*’ from the same R package. Previous to RDAs and pRDAs analyses, principal components analysis (PCA) on gene expression, genomic and epigenomic matrices were performed. Principal component (PCs) factors were used as the multivariate measure of gene expression, genomic and epigenomic variation. Only factors explaining at least 2.0% of the variation were kept. Selection of the best genomic and epigenomic PCs explaining the variance in gene expression was performed with backward selection, using the function ‘*ordistep*’ of *Vegan*. The selected PCs and associated variance are detailed in the TableS6. This procedure was repeated for comparisons involving limnetic and benthic species across continents and between limnetic and benthic species within each continent.

## DATA ACCESSIBILITY

Raw RNAseq sequence data are available through the NCBI sequence read archive (SRA) database under accession SRP136771. Raw WGBS sequence data are available through the NCBI sequence read archive (SRA) database under the BioProject PRJNA559821. The pipeline used for the whole genome bisulphite sequencing is available at: https://github.com/crougeux/wgbs_methylation_calling.

## AUTHOR CONTRIBUTIONS

C.R. and L.B. designed the study. CR performed bench work for DNA and RNA, treated the raw data, analysed the data for RNAseq and WGBS. CR and ML analysed the data. CR was the lead in writing of the manuscript, which also involved ML, P-A.G. and L.B. All authors contributed to writing and critically commented the final version of the manuscript.

## ACKNOWLEDGEMENTS

We would like to thank Anne-Marie Dion-Côté for insightful discussions around epigenomics in whitefish, as well as Jérémy Le Luyer for advices and exchanges about methodological approaches. We thank Kim Praebel, Shripathi Bhat and the Freshwater ecology group at UiT (Tromso) for providing samples from Norway, and Ole Seehausen for providing samples from Switzerland. This research was supported by a Discovery Research grant from the Natural Sciences and Engineering Research Council of Canada (NSERC) to L.B. L.B also holds the Canadian Research Chair in genomics and conservation of aquatic resources, which funded the research infrastructure for this project.

