## Supplementary for "The role of genomic vs. epigenomic variation in shaping patterns of convergent transcriptomic variation across continents in a young species complex"

**S1 Fig. Differential gene expression between limnetic and benthic whitefish induced by DNA methylation.** Plot showing the level of gene expression differentiation (Log2 fold change) as a function of the difference in methylation level between the limnetic and benthic species (ΔmCG Limnetic-Benthic) for genes associated with a convergent DMR. Each dot corresponds to a gene. Genes with negative ΔmCG have hypo-methylated DMRs in limnetic species, whereas genes with positive ΔmCG correspond to genes with hyper-methylated DMRs in the limnetic species. Both categories are respectively associated with an overexpression (black dots with a Log2 fold change > 0) and a repression (black dot with a Log2 fold change < 0) of gene expression and showed difference in degrees of gene expression levels ($P < 0.001$), while grey dots showed opposite patterns.

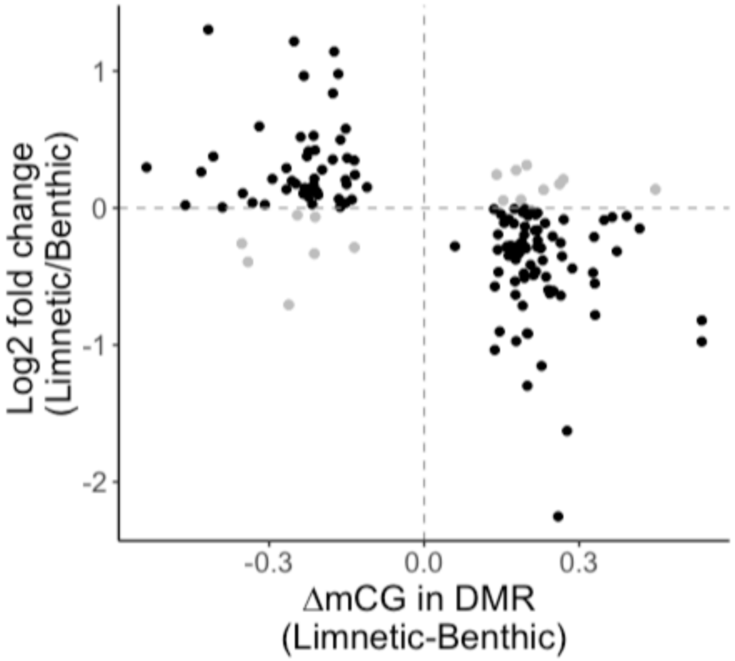

**S2 Fig.** Relationship showing that the percentage of repressed genes is higher when statistical significance for DMRs is higher.

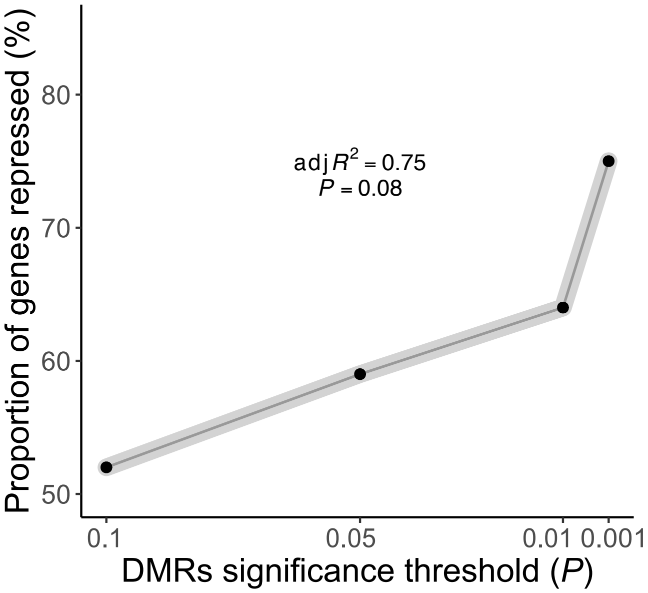

**S3 Fig. Distribution of inferred eQTL p-values.** QQ-plot distribution of the p-value of all tested eQTLs. Figure obtained directly from Matrix eQTL.

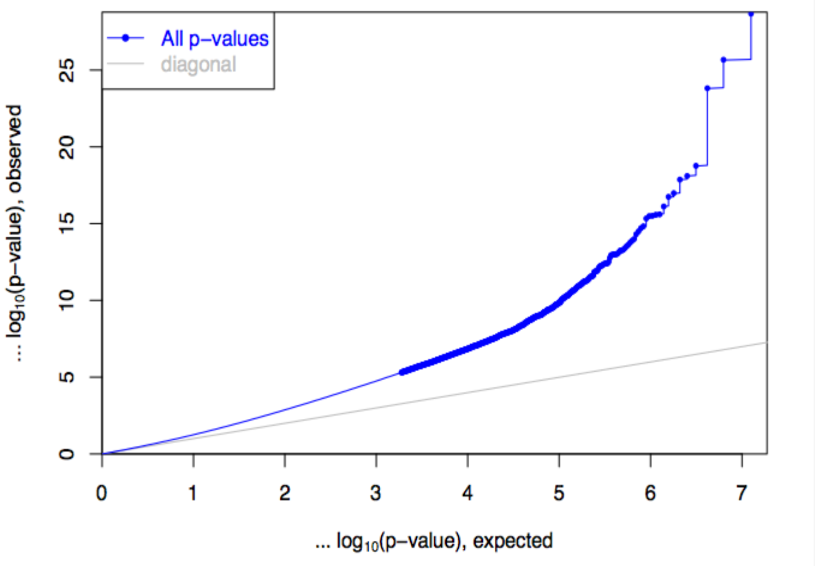

**S4 Fig. Differential gene expression between limnetic and benthic whitefish induced by DNA methylation (DMR) and cis-eQTL.** Boxplot showing the level of gene expression differentiation (Log2 fold change) between the limnetic and benthic species for genes associated with a parallel DMR, compared to genes associated with a *cis*-eQTL. Level of gene expression differentiation was higher in over-expressed genes in the limnetic species relative to the benthic species when harbouring a *cis*-eQTL (light-green box) compared to genes harbouring a DMR (light-blue box, $P < 0.001$). The same pattern emerged for under-expressed genes in the limnetic species relative to the benthic species in presence of DMR (purple box) compared to genes with eQTL (red box, $P = 0.012$).

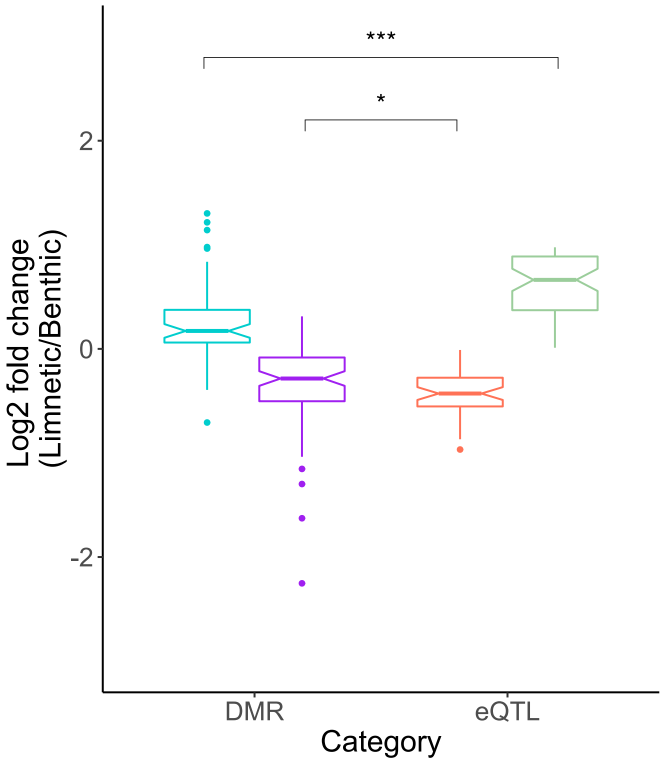

**S5 Fig. DMR characteristics depending on their statistical significance threshold.** A) DMRs had shorter length and less variation in length when identified with more stringent thresholds. B) Weak differences in the amount of CpGs per DMR despite lower length suggest a higher CpGs density in more significant DMRs. C) More significant DMRs showed higher methylation differences between limnetic and benthic species.

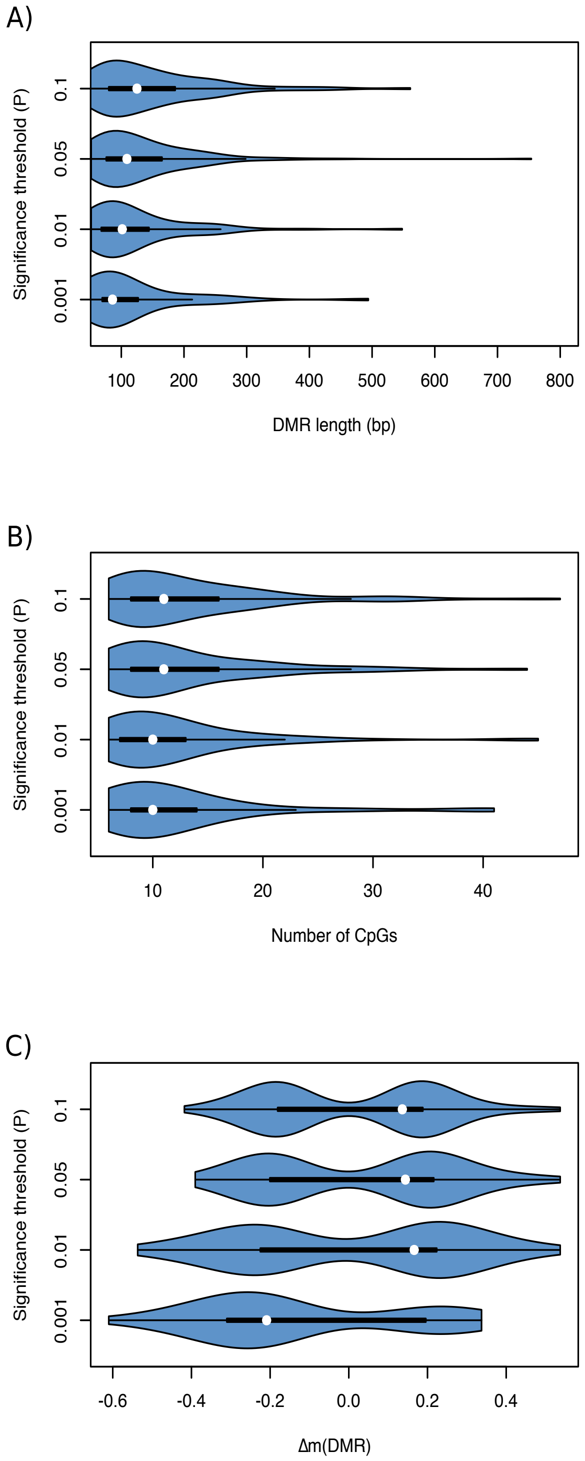

### Supplementary tables

#### Table S1: Number of filtered reads obtained per individual for transcriptomic data.

| Number of filtered reads | Library ID | Individual ID | Species |
|---|---|---|---|
| 25803019 | LIB58 | CD17 | Limnetic |
| 32293410 | LIB70 | CD18 | Limnetic |
| 27755684 | LIB52 | CD19 | Limnetic |
| 24577489 | LIB47 | CD20 | Limnetic |
| 19650600 | LIB43 | CD21 | Limnetic |
| 26670427 | LIB53 | CD22 | Limnetic |
| 24265292 | LIB16 | CN10 | Benthic |
| 19295658 | LIB11 | CN11 | Benthic |
| 25351805 | LIB37 | CN12 | Benthic |
| 18676563 | LIB60 | CN5 | Benthic |
| 26358596 | LIB6 | CN6 | Benthic |
| 31286322 | LIB68 | CN7 | Benthic |
| 25833236 | LIB30 | ID1 | Limnetic |
| 16896683 | LIB5 | ID2 | Limnetic |
| 22170784 | LIB19 | ID3 | Limnetic |
| 24150716 | LIB39 | ID4 | Limnetic |
| 26628586 | LIB13 | ID7 | Limnetic |
| 21907482 | LIB51 | ID9 | Limnetic |
| 19875693 | LIB71 | IN10 | Benthic |
| 20817338 | LIB25 | IN5 | Benthic |
| 20110167 | LIB38 | IN6 | Benthic |
| 15224135 | LIB44 | IN7 | Benthic |
| 16715432 | LIB21 | IN8 | Benthic |
| 23292157 | LIB31 | IN9 | Benthic |
| 34438149 | LIB66 | LF107 | Limnetic |
| 29159603 | LIB61 | LF108 | Limnetic |
| 29818408 | LIB18 | LF109 | Limnetic |
| 40665197 | LIB63 | LF111 | Limnetic |
| 21850810 | LIB2 | LF112 | Limnetic |
| 32821823 | LIB33 | LF113 | Limnetic |
| 23452258 | LIB54 | LF35 | Benthic |
| 22169931 | LIB9 | LF36 | Benthic |
| 21266681 | LIB48 | LF37 | Benthic |
| 23818507 | LIB12 | LF38 | Benthic |
| 27242770 | LIB56 | LF39 | Benthic |

| | | | |
|---|---|---|---|
| 22394148 | LIB27 | LF41 | Benthic |
| 18931029 | LIB41 | Z11 | Benthic |
| 27811520 | LIB28 | Z12 | Benthic |
| 14212670 | LIB34 | Z13 | Benthic |
| 18186632 | LIB29 | Z19 | Benthic |
| 21220544 | LIB49 | Z20 | Benthic |
| 20525864 | LIB17 | Z23 | Limnetic |
| 20566388 | LIB22 | Z27 | Limnetic |
| 29809348 | LIB40 | Z29 | Limnetic |
| 19688042 | LIB10 | Z30 | Limnetic |
| 19797211 | LIB64 | Z31 | Limnetic |
| 9001555 | LIB69 | Z37 | Limnetic |
| 26602363 | LIB4 | Z15 | Benthic |

**Table S2: Details of parallel DEGs between limnetic and benthic species, annotated as TEs.**

| Transcript | TE_name | TE_type | Superfamilly |
|---|---|---|---|
| TRINITY_DN53008_c5_g6_i3 | Rex1-21_DRe | REX | non-LTR |
| TRINITY_DN53328_c9_g1_i2 | TE-X-12_DR | undefined | transposon |
| TRINITY_DN55101_c2_g5_i1 | Crack-1_DR | L2 | non-LTR |
| TRINITY_DN78678_c0_g1_i1 | Copia-13E_DR-I | Copia | LTR |

**Table S3: Gene ontology analysis results for significant enrichment.** Significant GO term ($P$<0.05) from differential gene expression analyses between limnetic and benthic species.

| GO_ID | Biological level | | Biological function | FDR |
|---|---|---|---|---|
| GO:0006807 | BP | p | nitrogen compound metabolic process | 0.000597 |
| GO:0007165 | BP | p | signal transduction | 0.00151 |
| GO:0044237 | BP | p | cellular metabolic process | 0.00189 |
| GO:0044238 | BP | p | primary metabolic process | 0.00563 |
| GO:0071704 | BP | p | organic substance metabolic process | 0.00792 |
| GO:0065007 | BP | p | biological regulation | 0.00962 |
| GO:0050789 | BP | p | regulation of biological process | 0.0115 |
| GO:0009058 | BP | p | biosynthetic process | 0.0116 |
| GO:0032259 | BP | e | methylation | 0.0125 |
| GO:0008152 | BP | p | metabolic process | 0.0167 |
| GO:0051716 | BP | p | cellular response to stimulus | 0.0436 |
| GO:0032991 | CC | p | macromolecular complex | 0.000102 |
| GO:0043234 | CC | p | protein complex | 0.000716 |
| GO:0099080 | CC | e | supramolecular complex | 0.00204 |
| GO:0099081 | CC | e | supramolecular polymer | 0.00204 |
| GO:1990015 | CC | e | ensheathing process | 0.0048 |
| GO:0098576 | CC | e | lumenal side of membrane | 0.0143 |
| GO:0097223 | CC | e | sperm part | 0.022 |
| GO:0098948 | CC | e | intrinsic component of postsynaptic specialization membrane | 0.0284 |
| GO:0030672 | CC | e | synaptic vesicle membrane | 0.0348 |
| GO:0035749 | CC | e | myelin sheath adaxonal region | 0.0424 |
| GO:0098936 | CC | e | intrinsic component of postsynaptic membrane | 0.047 |
| GO:0016491 | MF | e | oxidoreductase activity | 0.0031 |
| GO:0003823 | MF | e | antigen binding | 0.0215 |
| GO:0004791 | MF | e | thioredoxin-disulfide reductase activity | 0.0331 |
| GO:0048037 | MF | e | cofactor binding | 0.043 |
| GO:0016209 | MF | e | antioxidant activity | 0.0447 |

**Table S4: Sequencing efficiency and coverage per WGBS library.** The identification of samples (C: Cliff Lake, I: Indian Lake, Lf: Langfjordvatn Lake and Z: Zurich lake; and L; Limnetic and B: benthic), the number of reads generated per library (nReads) and the individual coverage associated.

| ID | nReads | Coverage (X) |
|---|---|---|
| CL17 | 73428106 | 8.81137272 |
| CL18 | 128872591 | 15.46471092 |
| CL19 | 120073762 | 14.40885144 |
| CL20 | 114373473 | 13.72481676 |
| CL21 | 126932751 | 15.23193012 |
| CL22 | 128312915 | 15.3975498 |
| CB10 | 89843255 | 10.7811906 |
| CB11 | 101163627 | 12.13963524 |
| CB12 | 125040522 | 15.00486264 |
| CB5 | 110977507 | 13.31730084 |
| CB6 | 110086107 | 13.21033284 |
| CB7 | 136320804 | 16.35849648 |
| IL1 | 138579146 | 16.62949752 |
| IL2 | 111383287 | 13.36599444 |
| IL3 | 95107625 | 11.412915 |
| IL4 | 107098869 | 12.85186428 |
| IL7 | 103559153 | 12.42709836 |
| IL9 | 85605173 | 10.27262076 |
| IB10 | 101460525 | 12.175263 |
| IB5 | 100842607 | 12.10111284 |
| IB6 | 83510961 | 10.02131532 |
| IB7 | 106055635 | 12.7266762 |
| IB8 | 117124294 | 14.05491528 |
| IB9 | 119580386 | 14.34964632 |
| LfL107 | 113790085 | 13.6548102 |
| LfL108 | 124151997 | 14.89823964 |
| LfL109 | 110672336 | 13.28068032 |
| LfL111 | 116574416 | 13.98892992 |
| LfL112 | 132113031 | 15.85356372 |
| LfL113 | 105026245 | 12.6031494 |
| LfB35 | 110189564 | 13.22274768 |
| LfB36 | 114091761 | 13.69101132 |
| LfB37 | 121404423 | 14.56853076 |
| LfB38 | 122307506 | 14.67690072 |
| LfB39 | 127378901 | 15.28546812 |

| | | |
|---|---|---|
| LfB41 | 113703088 | 13.64437056 |
| ZL23 | 85282287 | 10.23387444 |
| ZL27 | 117595494 | 14.11145928 |
| ZL29 | 120073762 | 14.40885144 |
| ZL30 | 114373473 | 13.72481676 |
| ZL31 | 126932751 | 15.23193012 |
| ZL37 | 128312915 | 15.3975498 |
| ZB11 | 117595494 | 14.11145928 |
| ZB12 | 120073762 | 14.40885144 |
| ZB13 | 114373473 | 13.72481676 |
| ZB15 | 126932751 | 15.23193012 |
| ZB19 | 128312915 | 15.3975498 |
| ZB20 | 113790085 | 13.6548102 |

**Table S5: Summary table of the number of CpGs (raw and filtered) among population, and of DMLs and DMRs between species.**s. The number of DMLs and DMRs reported were identified when comparing the limnetic species to the benthic species, within lakes and across all populations.

| Population | n(CpGs) raw | n(CpGs) filtered | sd | DMLs | DMRs |
|---|---|---|---|---|---|
| Cliff L. | 1526526 | 864143 | 399951 | 21843 | 748 |
| Cliff B. | 1560891 | 995605 | 64301 | | |
| Indian L. | 1555263 | 1001147 | 95639 | 15878 | 532 |
| Indian B. | 1548708 | 1012432 | 39255 | | |
| Langfjordvatn L. | 1484524 | 914369 | 78472 | 16593 | 575 |
| Langfjordvatn B. | 1470490 | 883485 | 89648 | | |
| Zurich L. | 1523043 | 952037 | 75774 | 17730 | 619 |
| Zurich B. | 1514521 | 975403 | 74482 | | |
| All populations | 1522996 | 949828 | 114690 | 9819 | 525 |

**Table S6: Gene ontology analysis results for significant enrichment.** Significant GO term (*P*<0.05) from differentially methylated regions (DMRs) between all limnetic and all benthic whitefish across continents.

| GO_ID | Biological level | | Biological function | FDR |
|---|---|---|---|---|
| GO:0048856 | BP | e | anatomical structure development | 0.00161 |
| GO:0032502 | BP | e | developmental process | 0.00301 |
| GO:0044699 | BP | e | single-organism process | 0.00543 |
| GO:0009987 | BP | e | cellular process | 0.00588 |
| GO:0044767 | BP | e | single-organism developmental process | 0.00665 |
| GO:0022402 | BP | p | cell cycle process | 0.0159 |
| GO:0044464 | CC | e | cell part | 3.10E-06 |
| GO:0044424 | CC | e | intracellular part | 0.000128 |
| GO:0044422 | CC | e | organelle part | 0.000341 |
| GO:0044446 | CC | e | intracellular organelle part | 0.00216 |
| GO:0031090 | CC | e | organelle membrane | 0.00222 |
| GO:0044425 | CC | e | membrane part | 0.00929 |
| GO:0005622 | CC | e | intracellular | 0.0183 |
| GO:0044326 | CC | e | dendritic spine neck | 0.0223 |
| GO:1990794 | CC | e | basolateral part of cell | 0.0267 |
| GO:0098590 | CC | e | plasma membrane region | 0.0303 |
| GO:0044456 | CC | e | synapse part | 0.0318 |
| GO:0043226 | CC | e | organelle | 0.0374 |
| GO:0097458 | CC | e | neuron part | 0.0378 |
| GO:0060171 | CC | e | stereocilium membrane | 0.0397 |
| GO:0045281 | CC | e | succinate dehydrogenase complex | 0.0397 |
| GO:0044420 | CC | e | extracellular matrix component | 0.0446 |
| GO:0005488 | MF | e | binding | 2.98E-05 |
| GO:0043167 | MF | e | ion binding | 0.000644 |
| GO:0016491 | MF | e | oxidoreductase activity | 0.00179 |
| GO:0048037 | MF | e | cofactor binding | 0.00928 |
| GO:0051540 | MF | e | metal cluster binding | 0.011 |
| GO:0005515 | MF | e | protein binding | 0.0143 |
| GO:0050436 | MF | e | microfibril binding | 0.0178 |
| GO:0003824 | MF | e | catalytic activity | 0.0206 |
| GO:0005201 | MF | e | extracellular matrix structural constituent | 0.0356 |

**Table S7: Gene ontology analysis results for significant enrichment.** Significant GO term (*P*<0.05) from differentially methylated regions (DMRs) between all limnetic and all benthic whitefish across continents.

| Descriptive parameters | **All dataset** | **America** | **Europe** |
|---|---|---|---|
| **n of PCs axes with variance > 2%** | 7 | 16 | 12 |
| **Transcriptomic variance explained (%)** | 64.7 | 94.6 | 83.4 |
| **Backward selection - genomic (axis IDs)** | 1;2;11;14 | 1;2;3 | 1;10;13;22 |
| **Genomic variance explained (%)** | 40.9 | 33 | 29.3 |
| **Backward selection - epigenomic (axis IDs)** | 1;2 | 1;2;4;9;19 | 1;17;19 |
| **Epigenomic variance explained (%)** | 25.6 | 29.6 | 16.1 |
